# Plant Growth-Promoting Activity of Beta-Propeller Protein YxaL Secreted from *Bacillus velezensis* Strain GH1-13

**DOI:** 10.1101/471185

**Authors:** Yong-Hak Kim, Yunhee Choi, Yu Yeong Oh, Nam-Chul Ha, Jaekyeong Song

## Abstract

YxaL is conserved within *Bacillus subtilis* species complex associated with plant and soil. The mature protein YxaL contains a repeated beta-propeller domain, but the subcellular location and function of YxaL has not been determined. The gene encoding the mature YxaL protein was PCR amplified from genomic DNA of *B. velezensis* strain GH1-13 and used for recombinant protein production. A rabbit polyclonal antibody against the purified YxaL was generated and used for western blotting to determine the constitutive expression and secretion of YxaL, which exhibited a half-life of 1.6 h in the culture medium of strain GH1-13. In this study, we show that seed treatments of *Arabidopsis thaliana* and rice (*Oryza sativa L*) with less than 1 mg L^−1^ of purified YxaL in a soaking solution were effective at improving the root growth of plants. The seedlings of the treated *Arabidopsis* seeds markedly increased transcription of a 1-aminocyclopropane-1-carboxylate synthetase marker gene (ACS 11) but reduced expression of auxin-and abscisic acid-responsive marker genes (IAA1, GH3.3, and ABF4), especially when provided exogenous auxin. The horticulture experiments showed that pepper (*Capsicum annum*) seeds treated with 1 mg L^−1^ YxaL in soaking solution increased shoot growth and improved tolerance to drought stress. We hypothesize that YxaL secreted from plant growth-promoting *Bacillus* cells has a significant impact on plant roots, with the potential of improving plant growth and tolerance against stress.

## Introduction

The protein YxaL has been observed to interact with the DNA helicase PcrA in *Bacillus subtilis* [1]. This interaction can enhance the processivity of PcrA *in vitro,* but it is difficult to observe because the amino acid sequence of YxaL (formerly named YxaK) contains a signal peptide (1-44 amino acid residues) at the *N*-terminus for translocation across the cytoplasmic membrane [2]. Thus, it is difficult to deduce the site of subcellular interaction between the extracellular protein YxaL and the intracellular helicase PcrA (or DNA). After removing the *N*-terminal signal peptide, the mature YxaL protein (45-415 a.a.) is predicted to contain a repeated pyrrolo-quinoline quinone (PQQ) domain that forms a beta-propeller structure [1]. Beta-propeller proteins have diverse functions with different blade numbers [3]. Based on the sequence homology and structural similarity, it is postulated that beta-propeller homologs with different structure architectures of the blades may have originated from one ancestral blade, most likely that of a PQQ motif beta-propeller [4]. This beta-propeller domain is ubiquitous in diverse proteins with a similar beta-propeller fold observed in methanol dehydrogenase, which uses PQQ as cofactor [5]. In methanol dehydrogenases of methylotrophic bacteria, adjacent cysteine residues in active sites can form disulphide bridges for electron transfer reactions [6, 7]. In contrast, YxaL homologs in *Bacillus* species have no conserved cysteine residue, which suggest that their functions differ from that of PQQ-containing enzymes.

*B. velezensis* strain GH1-13 was isolated from rice paddy soil in Korea and can promote plant growth and suppress several pathogens [8]. The chromosome sequence of strain GH1-13 (GenBank accession number CP019040.1) contains highly homologous genes responsible for the biosynthesis of some plant hormones and secondary metabolites, such as indole-3-acetic acid, 2,3-butanediol, non-ribosomal lipopeptides, and polyketide antimicrobials, which are believed to be more proficient for rhizosphere colonisation and pathogen control than other members of the *B. subtilis* group [9–11]. In this paper, comparative gene analysis showed that *B. velezensis* strain GH1-13 contains a highly conserved sequence of the *yxaL* gene in the operational group of *B. amyloliquefaciens-velezensis-siamensis* within the *B. subtilis* species complex [12]. This gene is located in an operon composed of two genes, which were designated *yxaJL* in a previous study [13], but its subcellular location and function has not been defined.

The goal of this study was to investigate the localization and effect of YxaL on plant growth. The gene coding for the mature portion of YxaL (45-415 a.a.) was amplified from the genomic DNA of strain GH1-13 by PCR, and then cloned and transformed into *E. coli* to overproduce the protein. The purified protein was utilized as antigen to generate a rabbit polyclonal antibody for detection of YxaL by western blotting and applied for seed treatments of *Arabidopsis thaliana,* rice, and pepper in order to evaluate the effect on plant growth. We describe that YxaL was secreted from the *Bacillus* cells into the medium and had a positive effect on the growth of plant roots in the sub-nanomolar range.

## Materials and methods

### Strains and cultivation

*B. velezensis* strain GH1-13 was revived from frozen stocks in 50% glycerol at −80 °C and streaked onto tryptic soy agar (BD, Sparks, USA) plates. Single colonies were cultivated in tryptic soy broth (TSB) at 25 °C with aeration (180 rpm), and bacterial growth was assessed by measuring the optical density at 600 nm. For plant culture experiments, *A. thaliana,* rice (*Oryza sativa* L), and pepper (*Capsicum annum*) seeds were disinfected with 2% hypochlorite and 0.05% Triton-X for 10 min at room temperature (25 °C). Subsequently, the seeds were washed several times with sterile water, and then treated with various concentrations of purified YxaL (0 to 100 mg L^−1^) in a soaking solution for 2 h at room temperature. Treated seeds of *A. thaliana* and rice were planted on 0.5% NuSieve GTG agarose (FMC Bioproduct, Rockland, USA) plates containing 1% sucrose, and 0.05% MES buffer (pH 5.7) in 0.5× Murashige & Skoog (MS) medium [14], which was supplemented with or without indole-3-acetic acid (Sigma, St Louis, USA) at the final concentration of 0.5 μM. Pepper seeds were treated in the soaking solution with or without 1 mg L^−1^ YxaL for 2 h in room temperature, and dried for 30 min under air stream in a clean bench, and then were planted in each 5 replicates of plant tray (50 cases) containing horticultural soil with a 1:1 mixture of Baroker (Seoul Bio, Eumseong, Korea) and BM6 (Berger, Saint-Modeste QC, Canada).

### Cloning and expression of the *yxaL* gene

Genomic DNA of strain GH1-13 was extracted using a Wizard® Genomic DNA Purification kit (Promega, Madison, USA). A DNA fragment coding for the mature YxaL protein (45-415 a.a.) was amplified by PCR with the following primers: forward 5’-GGC<u>CCATGG</u>CGGAAACGGTATTTAAACAAAAT and reverse 5’-GGG<u>CTCGAG</u>TTATTTTTTTGCCCCGAATGCGA. The underlined *Nco*I and *XhoI* restriction sites are compatible with those in the plasmid pProEX-HTA with an *N*-His tag linked to a TEV protease cleavage site. After cloning the *yxaL* gene, the resulting plasmid pProEX-YxaL (*N*-His-TEV) was transformed into *E. coli* strain BL21 (DE3). The transformed cells were grown to an optical density of ~0.5 in 1 L of LB broth containing 100 mg ampicillin with aeration (180 rpm) at 37 °C, after which the recombinant YxaL protein was overexpressed by the addition of 0.2 mM IPTG for 1 h. Cells were harvested, treated with 1 mM mercaptoethanol and a protease cocktail (Roche Diagnostics, Indianapolis, USA), and disrupted by repeated ultrasonication in an ice-water bath.

Unless otherwise stated, protein purification was performed at 4 °C. After centrifugation at 21,000 ×*g* for 15 min, the supernatant was transferred to a new vessel, mixed with 20 mM imidazole and Ni NTA agarose (Qiagen, Hilden, Germany), and agitated on a rotary shaker for 1 h. The protein-bound agarose was loaded on a column, washed with 25 mM imidazole in 1×PBS (8 g NaCl, 0.2 g KCl, 1.44 g Na_2_HPO_4_, and 0.24 g KH_2_PO_4_ in 1000 mL water, pH 7.4), and the majority of YxaL (*N*-His-TEV) was eluted within the fractions containing 100 to 150 mM imidazole (Supplementary Figure 2a). After exchanging the buffer with 1×PBS using a G25 Sepharose desalting column, the purified protein was mixed with a His-tagged TEV protease at a ratio of 100:1 to remove the *N*-His-TEV site of the recombinant protein, as described previously [15]. The majority of the protein with the removed His tag was applied to an Ni NTA column and was eluted in the unbound fraction. The purified protein size and purity was determined by size exclusion chromatography and SDS PAGE.

### Western blotting

Purified YxaL was utilized as antigen to generate a rabbit polyclonal antibody for detecting YxaL by western blotting in subcellular compartments of strain GH1-13 during its growth in TSB medium. A purified anti-YxaL IgG was used as a primary antibody (dilution rate, 1:20,000), which was then detected using a secondary chicken anti-rabbit IgG antibody with horseradish peroxidase (HRP) conjugate (Abcam, Cambridge, UK) and a western blotting detection kit (Advansta, Menlo Park, USA). Chemiluminescent images were acquired using a ChemiDoc XRS image analyser (Bio-Rad, Hercules, USA), and images were manipulated using Molecular Dynamics ImageQuant version 5.2 (GE Healthcare, Waukesha, USA).

### Gene expression analysis

To determine the expression levels of the *yxaL* gene during different growth phases of *B. velezensis* strain GH1-13 and the phytohormone-responsive genes (IAA1, GH3.3, ACS 11, and ABF4) in 1-week-old cultured roots of YxaL-treated *Arabidopsis* seeds, RNA was extracted using Qiagen RNeasy mini kits. The cDNA was synthesized using a Qiagen QuantiTect^®^ Reverse Transcription Kit and quantitative PCR (qPCR) was performed using a Qiagen QuantiTect^®^ SYBR^®^ Green PCR Kit with a Roche LightCycler^®^ Nano instrument. The qPCR primers used are presented in S1 Table. Relative gene expression was calculated by the ΔΔCq method after normalization of the qPCR data to the 16S and 18S rRNA levels.

### Plant growth testing

To test effects of YxaL on plant growth, seeds of pepper (*Capsicum annuum*) treated with or without 1 mg L^−1^ YxaL were cultivated in greenhouse under normal watering conditions for 54 days to measure the length and biomass of shoots and roots. In order to examine the effect of YxaL on drought stress tolerance, the seedlings grown for 58 days in potted soil were deprived of water for 5 days, re-watered at intervals for 7 days and monitored for growth of leaves.

### Statistics

Experimental data from at least three independent replicates were reported as the means and standard deviation of the means. Significant differences between data were determined by analysis of variance (ANOVA) and t-tests, with differences considered to be significant at a P-value of less than 0.05.

## Results

### Evolutionary relationships of YxaL with beta-propeller domains in bacteria

A database search for YxaL homologs in bacteria showed that the amino acid sequence of YxaL obtained from *B. velezensis* strain GH1-13 is highly conserved in *Bacillus* species. The YxaL homologs are diverged into two types, tentatively named YxaL1 and YxaL2, which are present in distinct operational taxonomic groups of *B. amyloliquefaciens-siamensis-velezensis* and *B. halotolerans-nakamurai-tequilensis* (Fig 1 and S1 Fig). Among these strains, many containing YxaL1 have been observed to be primarily associated with plant and soil [12]. The beta-propeller protein BamB in *Mycobacteroides abscessus* subsp. *massiliense* is also included in the YxaL1 clade, indicating that the beta-propeller domain of BamB is commonly adopted in bacteria [4]. The consensus sequences of YxaL1 and YxaL2 were predicted to have the 8-blade propellers based on the PQQ motifs, AX(D/N)XXTG(D/E/K)XXW (Table 1; see S1 Datasets). YxaL1 and YxaL2 are distantly related to the other paralogs that are branched to from two beta-propeller domains (ingroup/outgroup) of *Escherichia coli* BamB and YncE, which include the only beta-propeller protein Rv1057 in *Mycobacterium tuberculosis.*

**Fig 1.**
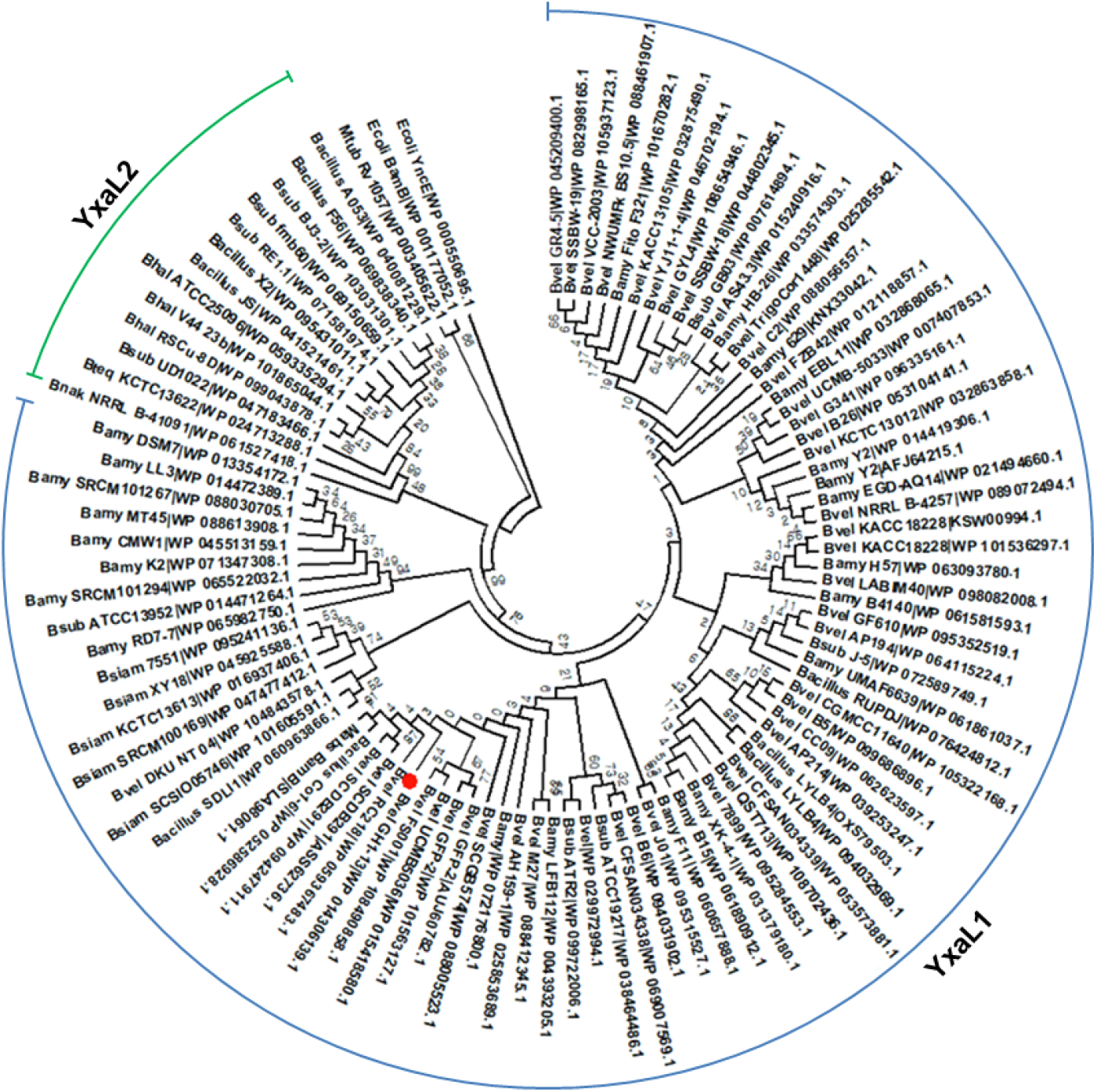
Phylogenetic analysis of homologous YxaL protein sequences. A minimum evolution tree was constructed using the Close-Neighbor-Interchange algorithm at a search level of 1, based on the Dayhoff model with 1000 bootstrap replications in MEGA 7. The Neighbor-Joining algorithm was used to generate the initial tree. The analysis involved 100 amino acid sequences. All ambiguous positions (gaps) generated from a multiple sequence alignment using Clustal Omega were removed for each sequence pair. There were a total of 630 positions in the final dataset and the evolutionary tree was drawn with branch lengths, as shown in S1 Fig. The position of YxaL obtained from *B. velezensis* strain GH1-13 is marked by a circle symbol before prefixing the strain name with the abbreviated taxonomic name. Abbreviations used: Bamy, *B. amyloliquefaciens*; Bhal, *B. halotolerans*; Bnak, *B. nakamurai*; Bsiam, *B. siamensis*; Bsub, *B. subtilis*; Bteq, *B. tequilensis*; Bvel, *B. velezensis*; Ecoli, *Escherichia coli*; Mabs, *Mycobacteroides abscessus* subsp. *massiliense*; and Mtub, *Mycobacterium tuberculosis.*

**Table 1.**
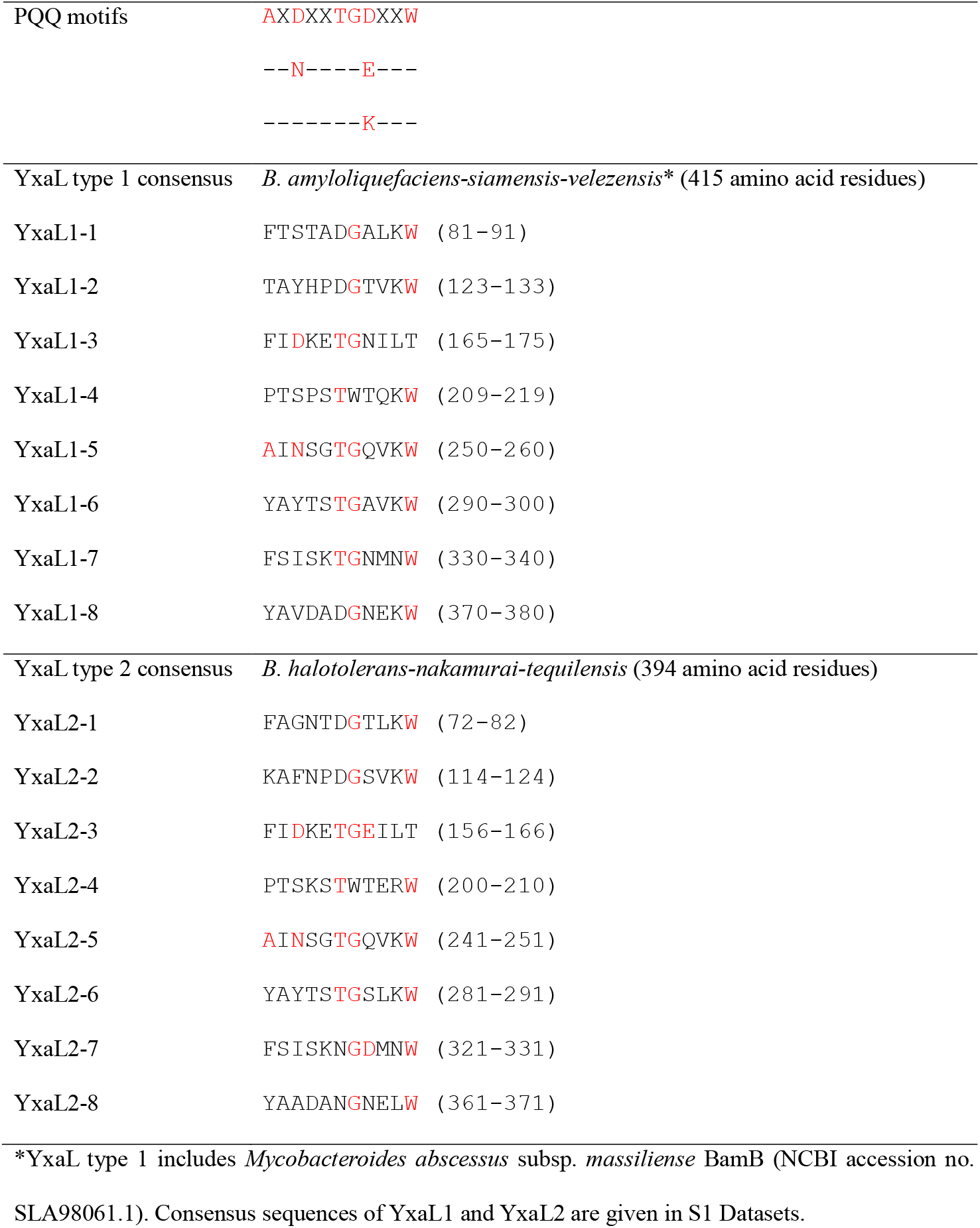
Beta-propeller motifs for two types of YxaL in the *Bacillus* species complex.

### Constitutive expression and secretion of YxaL in the medium of *B. velezensis* strain GH1-13

When cultivating strain GH1-13, constitutive expression of the *yxaL* gene was observed in the cell culture supernatants at different incubation times for early-exponential (5 h), late-exponential (8 h), early-stationary (12 h), and stationary (24 h) phases of growth (Fig 2A and B). The quantitative RT-PCR results for the *yxaL* gene transcript levels showed a similar pattern with the transcriptomic data in the NCBI BioProject PRJNA445855. Using genomic DNA of strain GH1-13 as a template, a DNA fragment coding for the mature YxaL protein (45-415 a.a.) was amplified by PCR and cloned into the plasmid pProEX-HTA with an *N*-His tag linked to the TEV protease cleavage site to develop a simple method of producing a recombinant protein (see S2 Fig). Briefly, the recombinant protein YxaL (*N*-His-TEV) was overproduced in *E. coli* and purified repeatedly using Ni NTA affinity chromatography before and after digestion with TEV protease (*N*-His) to remove the *N*-His-TEV site from the recombinant protein.

**Fig 2.**
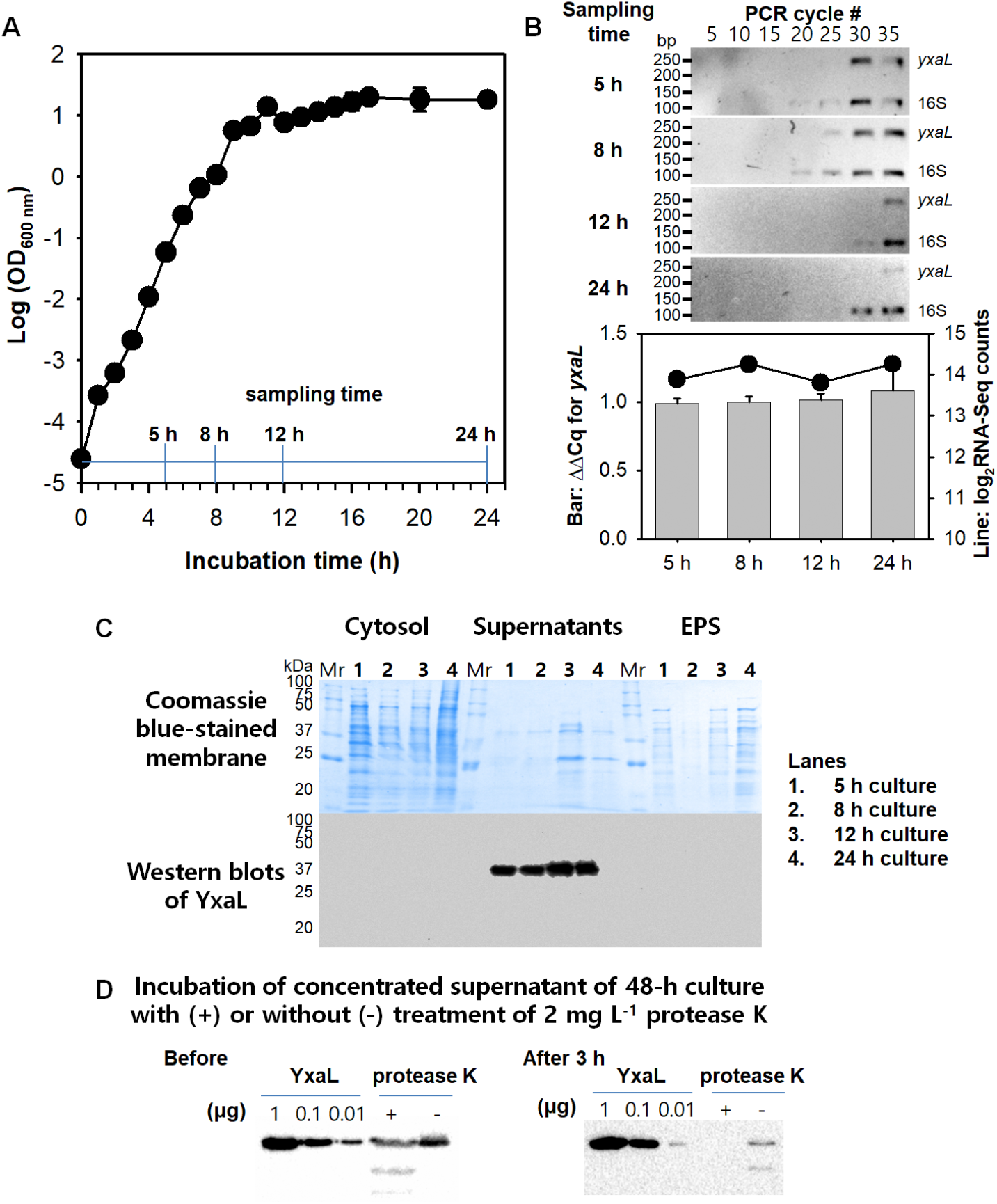
Constitutive expression and secretion of YxaL during cultivation of *B. velezensis* strain GH1-13. A: Growth curve of strain GH1-13 in tryptic soy broth with agitation (180 rpm) at room temperature (25 °C). Sampling time points for early-exponential (5 h), late-exponential (8 h), early-stationary (12 h), and stationary (24 h) phases of the curve are shown in the graph. B: Relative transcript (cDNA) levels of the *yxaL* gene to the 16S rRNA gene at the indicated sampling times for different growth phases of strain GH1-13. The upper panels show inverted gel images of semi-quantitative RT-PCR for expression levels of *yxaL* and 16S rRNA genes, which were evaluated every 5 cycles with a mixture of qPCR primers in S1 Table, and the lower graph shows the relative expression levels (ΔΔCq) of the *yxaL* gene to the 16S rRNA gene (bars) compared with the normalized log2-transformed values of the RNA-Seq counts obtained from the transcriptomic data deposited in the NCBI BioProject PRJNA445855. C: SDS PAGE and western blot analyses for determining the localization of YxaL in cytosol, supernatant, and extracellular matrix substance (EPS) fractions of the culture medium of strain GH1-13. The lower panels show the sensitivity and specificity of a polyclonal antibody generated using purified YxaL as the antigen (0.01 to 1 μg per lane) to determine the concentration and half-life of YxaL in (concentrated) supernatants of 48-h culture medium before and after a 3 h incubation with or without the addition of 2 mg L^−1^ protease K at room temperature.

Using this method, we achieved a high yield of 25 mg YxaL from 1 L LB medium cell culture, which was harvested after 1 h induction with 0.2 mM IPTG at the optical density (OD600) of 0. 5. Size exclusion chromatography showed that purified YxaL is a monomer with a molecular mass of 39,784. Using purified YxaL as an antigen, a rabbit polyclonal antibody was generated to determine the expression level and location of YxaL in the culture medium of strain GH1-13. The western blot results showed that YxaL was constitutively expressed and secreted from the cells, which produced approximately 100 μg L^−1^ YxaL in (concentrated) supernatants, with an observed half-life of 1.6 h (Fig 2C and D). These results confirm the previous proteome analysis of extracellular proteins in *B. subtilis* [2], and suggest that YxaL present in the *B. subtilis* species complex is constitutively expressed and excreted into the medium.

### Effects of YxaL on plant seedling growth

To investigate the effect of YxaL on plant growth, *A. thaliana* seeds were treated with various concentrations of YxaL in a soaking solution before planting on 0.5×MS medium solidified with 0.5% agarose, supplemented with or without 0.5 μM auxin. When the germination rate was evaluated 2 days after planting by counting germinated seeds under a microscope, the germination rate of YxaL-treated seeds was not different as compared to that of the corresponding untreated control group on either auxin-supplemented or unsupplemented media, although the germination of seeds on auxin-supplemented media displayed lower germination rates than those on unsupplemented media (Fig 3A). This is congruent with previous studies [16–18], suggesting that auxin plays a crucial role in seed dormancy. In contrast, YxaL-treated seeds planted in both auxin-supplemented and unsupplemented media exhibited similar lengths of 1-week cultured seedling roots, which were longer than those of the untreated seedling roots (Fig 3B). Furthermore, seedlings of treated seeds had markedly increased numbers of lateral and hair roots, compared to those of untreated seeds (Fig 3C). A similar effect of YxaL on the root growth and development was also observed for rice seedlings (see S3 Fig). From these results, the optimal concentration of YxaL in the soaking solution was determined to be 1 mg L^−1^ YxaL.

**Fig 3.**
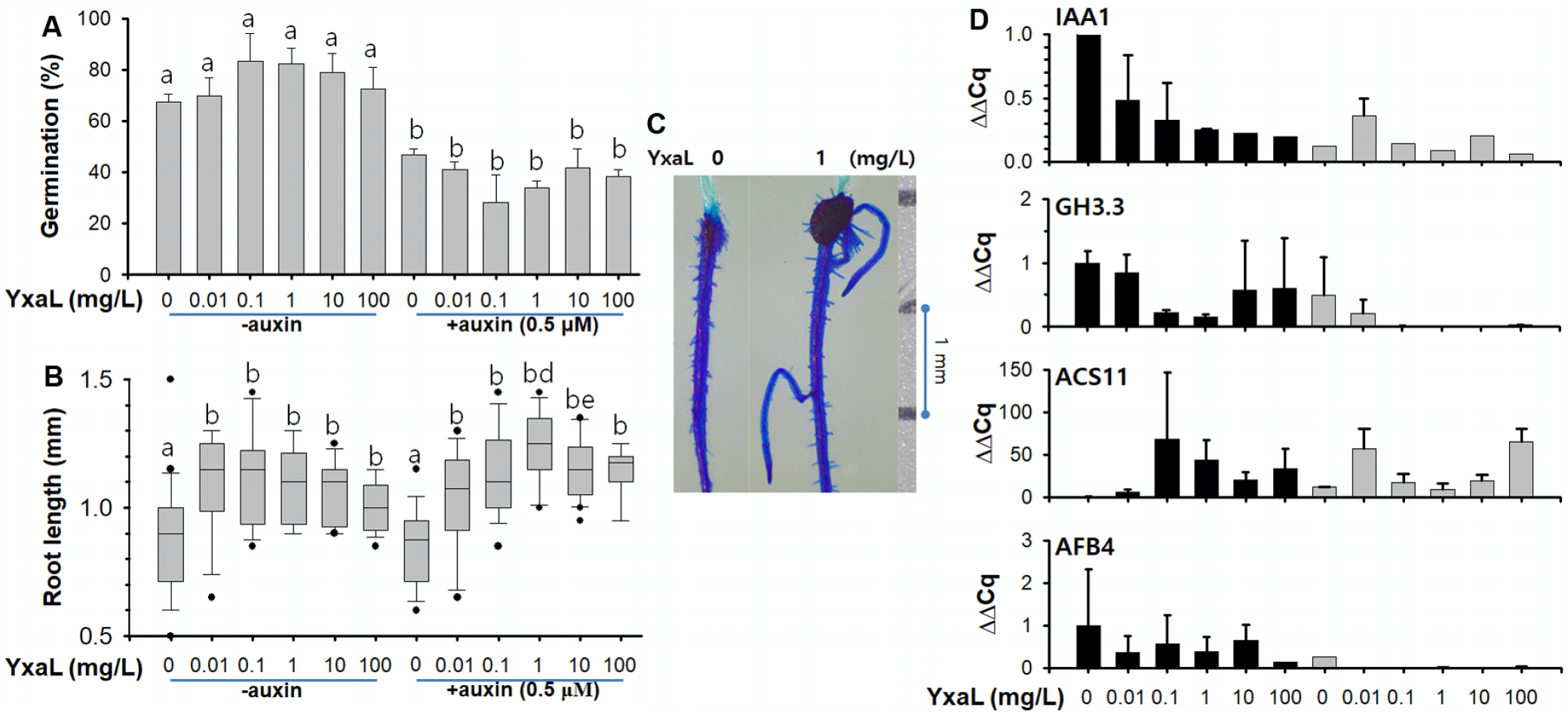
Differences in germination (A), root growth (B and C), and gene expression patterns (D) between YxaL-treated and untreated seeds of *A. thaliana* during seedling growth. **A**: Germination (%) of seeds (n > 100 for each group) treated with various concentrations of purified YxaL in soaking solutions and planted in auxin-supplemented and unsupplemented media. Germinated seeds were counted 2 days after planting. Compared to the untreated seeds (‘a’) in unsupplemented medium, significant differences determined by two-tailed t-tests with a *P*-value of less than 0.05 are indicated by the letter ‘b’ above the error bar. **B**: Root lengths of 1-week cultured seedlings (n > 30 for each group) in auxin-supplemented and unsupplemented media after seeds were treated with various concentrations of purified YxaL in the soaking solution. Significant differences determined by two-tailed t-tests with a P-value of less than 0.05 are shown above the box plot error bar with the letter ‘b’ to denote the comparison with the untreated seeds (‘a’) and followed with the letter ‘d’ or ‘e’ to denote comparisons between paired groups for seed treatment with 1 or 10 mg L^−1^ YxaL solution before planting in auxin-supplemented and unsupplemented media. **C**: Comparison of root architecture between 1-week seedlings of YxaL-treated and untreated seeds in unsupplemented media. The root area is stained with methylene blue and a scale bar with 1-mm intervals is shown at the right. D: Relative expression levels (ΔΔCq) of auxin-responsive protein (IAA1), indole-3-acetic acid amino synthetase (GH3.3), 1-aminocyclopropane-1-carboxylic acid (ACC) synthase (ACS11), and abscisic acid (ABA)-responsive element binding factor (AFB4) in 1-week-old seedling roots grown in auxin-supplemented and unsupplemented media after seed treatment with various concentrations of purified YxaL in the soaking solution. The results, which were obtained from triplicate culture experiments with a sample size (n) of greater than 30 for each group of 1-week-old roots, are reported as the means and standard deviation (error bar).

Using RNA extracts from 1-week-old *Arabidopsis* roots, RT-PCR was performed to assess changes in relative expression (ΔΔCq) of genes encoding auxin-responsive protein (IAA1), indole-3-acetic acid amino synthetase (GH3.3), 1-aminocyclopropane-1-carboxylic acid (ACC) synthase (ACS11), and abscisic acid (ABA)-responsive element binding factor (AFB4), which are responsive to auxin, ethylene, and ABA in the roots of *A. thaliana* at the early and later seedling stages [19]. The qPCR results showed that seed soaking in a low concentration of YxaL (10 to 100 μg L^−1^) exhibited markedly increased levels of ACS11 expression in the 1-week cultured roots by 34.5 fold within a range of 6.8 to 68 folds, compared to the untreated seedling roots, while they decreased expression of IAA1, GH3.3 and ABF4 (Fig 3D). These changes were similar to those observed in auxin-supplemented media. Moreover, when YxaL-treated seeds were cultivated in auxin-supplemented media, they exhibited stronger inhibition of GH3.3 and AFB4 gene expression than was observed for IAA1. The altered expression levels of the above four genes indicate that YxaL has a significant effect on other signaling pathways other than auxin for plant growth and development.

### YxaL testing for plant growth and drought stress tolerance

The observed root morphology of YxaL-treated *Arabidopsis* and rice indicated that YxaL might have a common effect on the root development and growth at a late stage regardless of the eudicot and monocot plants, grown under normal watering and waterlogging conditions. The root system architecture response has been proposed to account not only for the plant growth [20], but also for the functionality of plasticity in root development for tolerance [21]. We observed that, when YxaL-treated and untreated seeds of pepper (*Capsicum annuum*) were cultivated in potted soil under normal watering conditions for 54 days, YxaL-treated seeds had higher growth rate than the untreated seeds (Fig 4A). Stem heights and fresh weights of seedlings were significantly different between the two groups (two-tailed t-tests: *P* < 0.05), but no statistically significant difference between the weights of the seedling roots was noted (P = 0.42), though the weights of YxaL-treated seedling roots were slightly higher than those of the untreated seedling.

**Fig 4.**
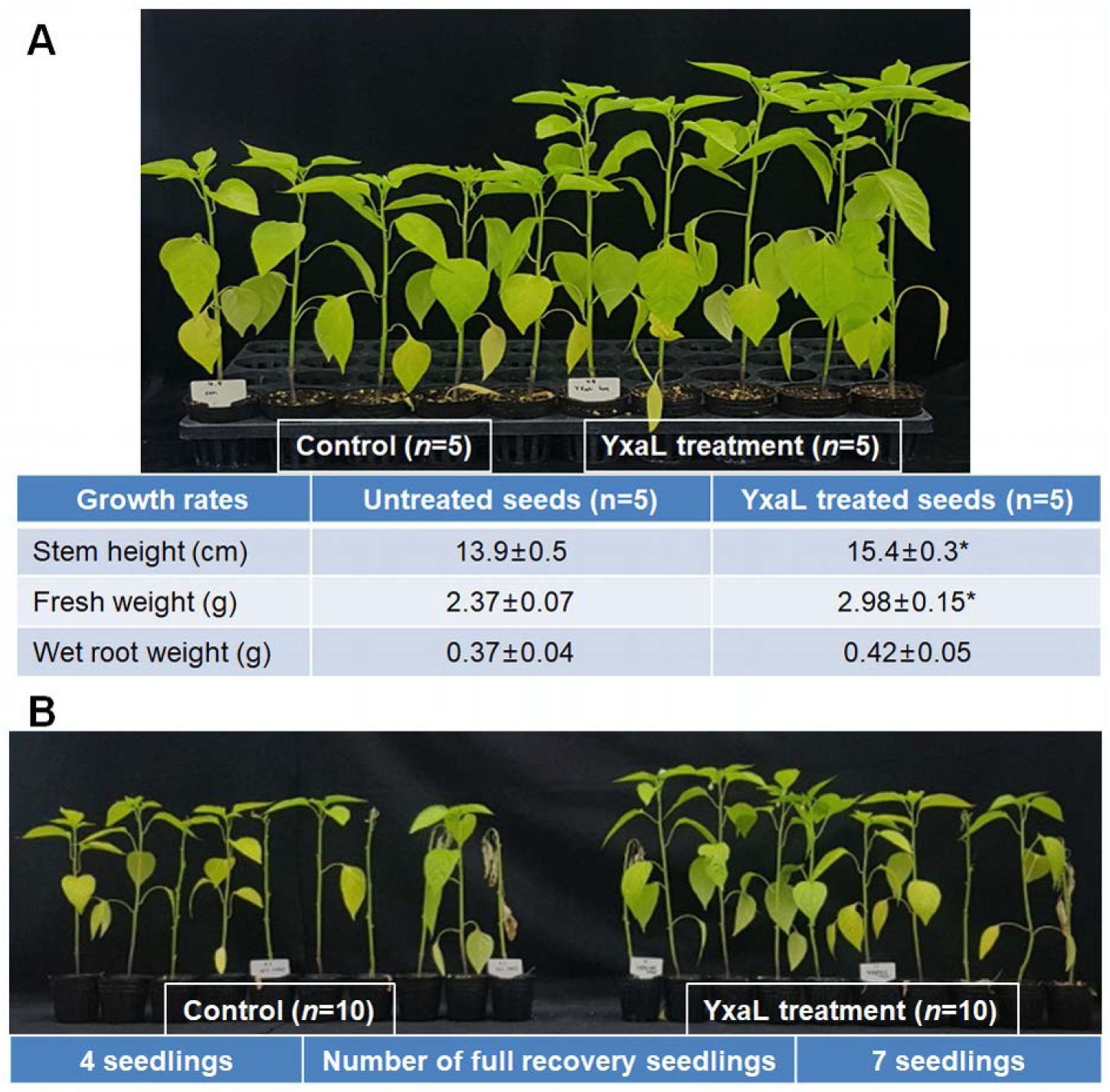
Effects of Yxal-treated pepper seeds on seedling growth and tolerance against drought stress. A: Shoot growth of pepper seedlings planted in potted soil after treatment of seeds in the soaking solution with or without 1 mg L^−1^ YxaL and grown in a greenhouse under normal watering conditions for 54 days. The means ± standard deviations of the measured data for stem height, fresh weight, and wet root weight were shown below the picture. Statistically significant differences between the data of the YxaL-treated and untreated groups were determined by two-tailed t-tests with the P-value of less than 0.05, as denoted by asterisks. B: Recovery of leaf status from drought stress-treated pepper seedlings of YxaL-treated and untreated seeds. After 58 days of culture in potted soil under normal watering conditions, drought stress was applied for 5 days, followed by re-watering for 7 days in order to count the number of the full recovery seedlings in each group of 10 seedlings, as noted below.

Furthermore, YxaL-treated seedlings exhibited improved tolerance to drought with greater potential than the untreated seedlings to recover the growth of leaves after re-watering (Fig 4B). After drought stress was applied for 5 days, followed by re-watering for 7 days, a full recovery of leaf status was observed in 7 seedlings of YxaL-treated seeds in a group of 10 seedlings, while at the same time only 4 seedlings of the untreated seeds in another group of 10 seedlings recovered full leaf turgor and resumed growth. These results indicate that YxaL-treated pepper seeds are effective in reducing the leaf damage caused by drought stress, together with alterations in root architecture during the seedling stage, and suggest that the development and architecture of roots from YxaL-treated seeds may have potential to increase plant growth and tolerance to drought stress.

## Discussion

We noted that the conserved sequences of YxaL in the plant growth-promoting *Bacillus* species complex are distantly related to the 8-blade beta-propeller protein BamB, which is essential in assembling outer membrane proteins in *E. coli* [22, 23]. This beta-propeller protein family may have diverse functions outside of the cell, as exemplified by the only beta-propeller protein Rv1057 in *M. tuberculosis,* which plays a role in the secretion of the major virulence factor ESAT-6 and contributes to cytotoxicity in infected macrophages [24]. The evolutionary tree shows that *E. coli* YncE is a distant outgroup with low sequence similarity to other proteins. Analyses of large-scale genome and outer membrane vesicle proteome data support that a highly immunogenic YncE is broadly expressed and secreted by different pathotypes of *E. coli* [25, 26]. Although secreted beta-propeller proteins in bacteria are largely unexplored, mounting data suggest that they may be involved in maintenance of the cell envelope, secretion, adhesion, and host immune responses.

In this study, the mature YxaL protein was deduced from the genomic DNA of *B. velezensis* strain GH1-13 and utilized for recombinant protein production and purification. Using the purified YxaL protein as an antigen, a polyclonal antibody was produced to determine the constitutive expression of secreted YxaL in supernatants by western blotting, the results of which were consistent with the transcriptomic and RT-PCR data. From the plant growth experiments, *A. thaliana* seeds soaked in a solution containing YxaL in the sub-nanomolar range (0.1 ~ 1 mg L^−1^) showed a significant improvement with respect to root growth. A previous study showed that the use of bovine serum albumin (BSA) as an organic N-source supplement produced similar effects on the root growth and development of *A. thaliana* at an optimal concentration of 1 g L^−1^ in the medium[27]. The effective concentration of BSA used in the medium was much higher than that of YxaL in a soaking solution which was useful to support plant growth-promoting activity for *A. thaliana* and rice by the seed soaking for 2 h before planting.

The RT-PCR results for relative gene expression of phytohormone-responsive genes (IAA1, GH3.3, ACS11, and AFB4) in 1-week-old *A. thaliana* roots demonstrated that YxaL has a significant effect on the upregulation of ACS11, which is responsible for the synthesis of ACC, the precursor of ethylene. However, the YxaL treatment decreased the expression of the other assayed genes (IAA1, GH3.3, and AFB4), which respond to auxin and ABA. It has been well established that the expression of ACS11 is induced in all cell types of the cell division zone in response to auxin and other stresses [28]. If the ACS11 gene is under the control of Aux/IAA-auxin response factors, it should be activated by the proteolytic degradation of the Aux/IAA transcriptional repressors in response to a specific signalling cue or by the binding of YxaL to dividing cells and tissues during seedling growth and not just during the germination stage [29]. The above gene expression pattern in the roots of 1-week-old *A. thaliana* seedlings, which germinated from seeds soaked in a solution containing YxaL, was somewhat similar to that seen in the auxin-supplemented media. It seems likely that exogenous auxin also enhances the expression of ACS11 in dividing cells [28], whereas it reduces germination rate in seeds, whether they are treated or not with YxaL. In contrast, the YxaL treatment of seeds did not affect germination and improved the root growth and development after germination, even in the presence of exogenous auxin. These events appeared to involve a pathway in which YxaL may interact with Aux/IAA-auxin response factors in a vast array of auxin signalling pathways, including activation of ACS11 during seedling growth, although the underlying mechanism is unknown.

ACC synthesis can help plants to not only synthesize ethylene, which antagonizes auxin and ABA to form branch roots [30], but also cope with stress by the induction of an ACC deaminase that can control endogenous levels of ACC and ethylene [31]. Plant-produced ACC can attract plant growth-promoting rhizobacteria with ACC deaminases, which can utilize ACC as organic N source, lowering ethylene production in plants and conferring tolerance to various stresses, such as flooding [32, 33], drought [34], salinity [35, 36], flower senescence [37], metal pollution [38], and pathogens [39]. The presence of an ACC deaminase in *Rhizobium leguminosarum* is likely to promote nodulation of pea plants by modulating ethylene levels in the plant roots during the early stages of nodule development [40]. Further studies remain to elucidate YxaL interactions with hormone response factors and metabolic enzymes, which may play pivotal roles in root growth and development by mediating interactions between plants and rhizobacteria.

## Conclusion

*B. velezensis* strain GH1-13 constitutively expressed and secreted a highly conserved beta-propeller protein YxaL in the taxonomic operational group of *B. amyloliquefaciens-velezensis-siamensis,* which is thought to be primarily associated with soil and plants. The mature protein YxaL overexpressed in *E. coli* was useful for treatment of plant seeds in the soaking solution with the optimal concentration of 1 mg L^−1^ to promote the development and growth of lateral and hair roots in different plant species. Horticulture experiments were performed to address the effect of YxaL treatment with relationship to alterations in the root system architecture which demonstrated that YxaL-treated pepper seeds had potential to increase plant yield and tolerance against drought stress. Our findings propose that YxaL is a new biomaterial with potential applications for promoting plant growth and improving tolerance against stress.

## Acknowledgements

This study was supported by a cooperative research programme for Agricultural Science & Technology Development (PJ012467062017), Rural Development Administration, Republic of Korea.

## Supporting Information

**S1 Datasets**. 97 homologous sequences of YxaL and 3 paralogous sequences of *Mycobacterium tuberculosis* Rv1057, *Escherichia coli* BamB and YncE were collected from the National Center for Biotechnology Information to analyse the phylogenetic trees (Fig 1 and S1 Fig) and to search the beta-propeller motif sequences (Table 1) in the consensus sequences of YxaL1 and YxaL2 generated by multiple sequence alignments.

**S1 Fig. Evolutionary relationship of YxaL homologs**.

The evolutionary history was inferred using the minimum evolution method in MEGA 7. The tree out of 46 minimum evolution trees (sum of branch length = 5.46420670) is shown, with branch lengths in the same units as those of the evolutionary distances used to infer the phylogenetic tree.

**S2 Fig. Recombinant protein production and purification of the mature YxaL protein using an Ni NTA agarose column.**

The recombinant protein YxaL with the *N*-His TEV cleavage site was purified by stepwise elution with high concentrations of imidazole (50 to 250 mM) and, after the #-His TEV cleavage site was removed from the recombinant protein YxaL by overnight digestion with a 1:100 ratio of a recombinant TEV protease (*N*-His), the mature protein YxaL was recovered in unbound fraction eluted by low concentrations of imidazole (20 to 25 mM) and the protein size and purity was determined by size exclusion chromatography and SDS PAGE.

**S3. Fig. Effects of soaking seeds with YxaL on the root growth and development of rice (*Oryza sativa* L.).**

After planting of rice seeds soaked with various concentrations of YxaL (0 to 100 mg L^−1^), the primary and hair root lengths were measured on 1 week after seedling emergence.

**S1 Table. Primers used in qPCR for determination of the expression levels of the *Bacillus yxaL* gene and *Arabidopsis* genes.**

See Fig 2 and Fig 3 for the relative expression levels of the *yxaL* gene in *Bacillus velezensis* strain GH1-13 and plant hormone-responsive marker genes (IAA1, GH3.3, AFB4, and ACS11) in *Arabidopsis thaliana* normalized to the respective 16S and 18S rRNA levels by the ΔΔCq method.

